# Single-cell analysis of colonic epithelium reveals unexpected shifts in cellular composition and molecular phenotype in treatment-naïve adult Crohn’s disease

**DOI:** 10.1101/2021.01.13.426602

**Authors:** Matt Kanke, Meaghan M. Kennedy, Sean Connelly, Matthew Schaner, Michael T. Shanahan, Elisabeth A. Wolber, Caroline Beasley, Grace Lian, Animesh Jain, Millie D. Long, Edward L. Barnes, Hans H. Herfarth, Kim L. Isaacs, Jonathan J. Hansen, Muneera Kapadia, José Gaston Guillem, Terrence S. Furey, Shehzad Z. Sheikh, Praveen Sethupathy

**Affiliations:** Department of Biomedical Sciences, Cornell University, Ithaca, NY, USA; Center for Vertebrate Genomics, Cornell University, Ithaca, NY, USA; Center for Gastrointestinal Biology and Disease, University of North Carolina at Chapel Hill, Chapel Hill, NC, USA; Department of Genetics, University of North Carolina at Chapel Hill, Chapel Hill, NC, USA; Curriculum in Bioinformatics and Computational Biology, University of North Carolina at Chapel Hill, Chapel Hill, NC, USA; Department of Surgery, University of North Carolina at Chapel Hill, Chapel Hill, NC, USA; Department of Biology, University of North Carolina at Chapel Hill, Chapel Hill, NC, USA

**Keywords:** Crohn’s disease, single-cell, epithelium, colonocyte, gene expression, ISC, EEC, SPIB

## Abstract

The intestinal epithelial barrier is comprised of a monolayer of specialized intestinal epithelial cells (IECs) that are critical in maintaining gut mucosal homeostasis. Dysfunction within various IEC fractions can increase intestinal permeability, resulting in a chronic and debilitating condition known as Crohn’s disease (CD). Defining the molecular changes in each IEC type in CD will contribute to an improved understanding of the pathogenic processes and the identification of potential therapeutic targets. Here we performed, for the first time at single-cell resolution, a direct comparison of the colonic epithelial cellular and molecular landscape between treatment-naïve adult CD and non-IBD control patients. Our analysis revealed that in CD patients there is a significant skew in the colonic epithelial cellular distribution away from canonical *LGR5*+ stem cells, located at the crypt-bottom, and toward one specific subtype of mature colonocytes, located at the crypt-top. Further analysis revealed unique changes to gene expression programs in every major cell type, including a previously undescribed suppression in CD of most enteroendocrine driver genes as well as L-cell markers including *GCG*. We also dissect a previously poorly understood *SPIB*+ cell cluster, revealing at least four sub-clusters that exhibit unique features. One of these *SPIB*+ sub-clusters expresses crypt-top colonocyte markers and is significantly up-regulated in CD, whereas another sub-cluster strongly expresses and stains positive for lysozyme (albeit no other canonical Paneth cell marker), which surprisingly is greatly reduced in expression in CD. Finally, through integration with data from genome-wide association studies, we show that genes implicated in CD risk exhibit heretofore unknown cell-type specific patterns of aberrant expression in CD, providing unprecedented insight into the potential biological functions of these genes.

## Introduction

The colonic epithelium acts as an essential barrier between the luminal contents of the colon, including a diverse compendium of microbes, and the underlying lamina propria immune system. The epithelium is a heterogenous mix of distinct cell types with a wide range of specialized functions, including absorptive cells that transport nutrients and electrolytes (colonocytes), and secretory cells that emit factors such as mucins (goblet cells) and endocrine hormones (enteroendocrine cells). Stem cells at the base of the colonic crypt are responsible for the continual, rapid renewal of this epithelial layer. Each cell type is crucial for the maintenance of intestinal homeostasis and defects in any could contribute to the onset of inflammatory bowel disease (IBD).

IBD consists of two main disease types, ulcerative colitis (UC) and Crohn’s disease (CD), and is characterized by chronic intestinal inflammation that can lead to severe tissue damage and organ dysfunction. Despite recent advances, the etiology of IBD still remains largely unknown. Unrelenting inflammation is attributed to a complex interaction between genetic, luminal (microbial), and environmental factors that trigger an inappropriate mucosal immune response. Recent studies have begun to unravel the role of different colonic epithelial cell types in IBD using single-cell RNA sequencing (scRNA-seq) (Elmentaite et al., 2020; Parikh et al., 2019; Smillie et al., 2019). While they have advanced our understanding, important challenges remain to be addressed. Notably, the patients included in the studies exhibit varying disease durations, and some have been treated with therapeutics, which are confounding variables. No scRNA-seq study to date has investigated treatment-naïve adult IBD. Moreover, the studies have focused mostly on patients with UC. No scRNA-seq study to date has performed a focused investigation of colonic epithelium in adult CD. Changes to the relative abundance and molecular character of different colonic epithelial cell types during CD pathogenesis is poorly understood and merits deeper examination.

To investigate this, we performed single-cell transcriptional profiling in epithelial cells from a cohort of treatment-naïve adult CD patients (n=3) and healthy controls (n=4). Our analysis reveals that the colonic epithelium of CD patients exhibits significant and unexpected shifts in cellular composition as well as cell-type specific transcriptional profiles, providing important clues about the early molecular events that promote the dysfunction of this critical tissue during CD pathogenesis.

## Results

### Single-cell analysis provides a high resolution picture of the colonic epithelium from adult non-IBD and Crohn’s patients

We harvested mucosal tissue from the ascending colon of treatment-naïve adult individuals with Crohn’s disease (CD, n=3) and non-IBD healthy controls (NIBD, n=4) (Table S1). We then enriched for epithelial cells, which were dissociated into single-cell suspensions (Methods). Samples were subjected to RNA sequencing using the 10X Chromium platform (Methods) and 13,039 cells remained after filtering (Methods).

We processed and analyzed the data (CD and NIBD together) using Cell Ranger and Seurat (Methods) and visualized the cell clusters using Uniform Manifold Approximation and Project (UMAP)(Becht et al., 2018). We identified 21 cell clusters (Figure S1A), 7 of which were determined to be different types of immune cells (Figure S1B), representing 21% of the total. We removed these cells and re-clustered the remaining 10,162 epithelial cells, resulting in 14 cell clusters. Based on the expression levels of previously annotated marker genes (Smillie et al., 2019), we assigned these clusters to 14 different colonic epithelial cell types (Figure 1A,B). These cell types include *LGR5*+ stem cells; *MUC2*+ immature and mature goblet cells; *CHGA*+ enteroendocrine cells (EECs); three categories of cycling cells including both G2-M-G1 and S-phase transit-amplifying (TA) cells, secretory progenitors (SPs), colonocyte progenitors; and at least four other cell clusters expressing known markers of colonocytes including *CA1*+ early and late colonocytes, *CEACAM7*+ colonocytes, and *SPIB*+ cells (Figure 1B,C). We found that each of these cell types express a unique set of markers (Methods, Figure 1D). Specifically, the *SPIB*+ cells are uniquely marked by *SPIB, NOTCH2* and *HES4*; the *CEACAM7*+ cells uniquely express several genes including one long, non-coding RNA (*LINC01133*) that has been implicated previously in cancer and in the regulation of the Wnt signaling pathway (Yang et al., 2018); and the *CA1*+ late colonocytes share *MALL* expression with *CEACAM7*+ cells but uniquely express *CA1* (Figure 1D).

**Figure 1:**
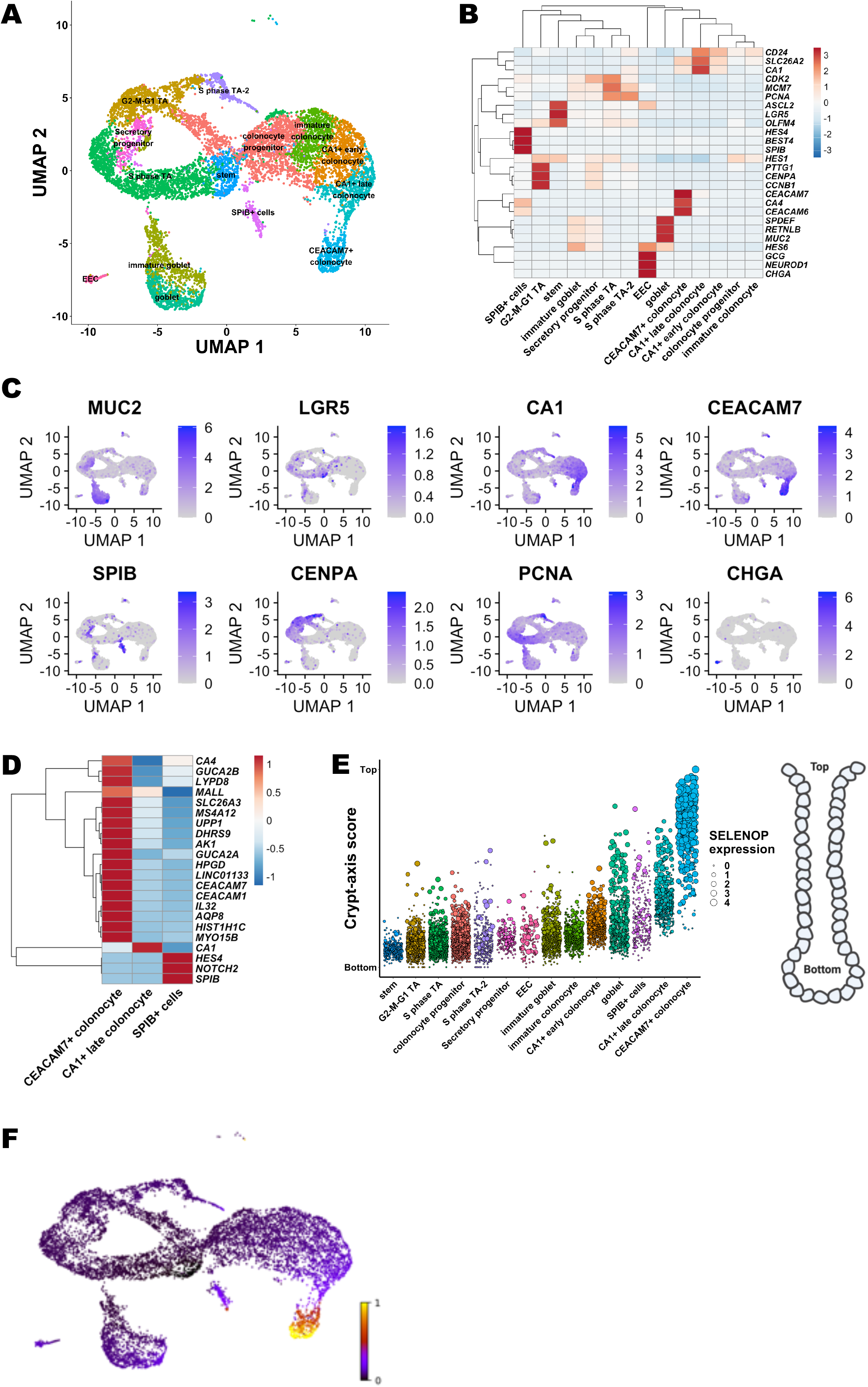
Single cell landscape in treatment-naïve Crohn’s disease and NIBD patients. (A) UMAP of epithelial cell clusters following assignment of cell type. (B) Normalized expression heatmap of genes known to be markers of various epithelial cell populations or found to be highly enriched in a cluster (rows) across epithelial clusters (columns) confirms cell type assignment. Normalized gene expression is scaled by row. (C) UMAP overlain with the normalized expression of cell type markers further confirms cluster assignment. MUC2 = goblet, LGR5 = stem, CA1 = CA1+ colonocytes, CEACAM7 = CEACAM7+ colonocytes, SPIB = SPIB+ cells, CENPA = G2-M-G1 TA, PCNA = S phase TA, CHGA = EEC. (D) Normalized expression heatmap of three clusters with a colonocytic signature (CA1+ late colonocytes, CEACAM7+ colonocytes, and SPIB+ cells) reveal expression of cluster-specific markers. Normalized expression is scaled by row. (E) Crypt-axis scores (low near crypt bottom, high near crypt top) of cells reveals expected location of clusters along the crypt axis. Clusters are arranged on x-axis by mean crypt-axis score (Methods). The size of the dot corresponds to level of expression of known marker of the crypt top, *SELENOP*. Diagram of colonic crypt shown at right. (F) UMAP overlain with the RNA velocity. RNA velocity is the ratio of spliced transcripts / unspliced transcripts. Scale is normalized to 1. NIBD = Non-inflammatory bowel disease, CD = Crohn’s disease, UMAP = Uniform Manifold Approximation and Projection, EEC = enteroendocrine cells.

Colonic epithelial cells from the crypt-bottom to the crypt-top represent a gradient of maturation, with stem/progenitor cells at the bottom and mature differentiated colonocytes at the top. Using a previously defined 15-gene signature (Parikh et al., 2019) we computed a crypt-axis score for every cell in each cluster (Figure 1E), which shows good correspondence between our cluster annotations and known cell-type positions within the crypt. RNA velocity analysis (La Manno et al., 2018) confirmed that the dataset represents the full maturational spectrum of the colonic epithelium (Figure 1F).

### The colonic epithelial cellular landscape is skewed toward a crypt-top signature in Crohn’s disease

Next we assessed CD and NIBD data separately to determine whether the distribution of cells along the crypt-axis, based on their crypt-axis score (Figure 1E), is altered in CD. We detected a significant shift in the density of cells toward the crypt top in CD relative to NIBD (Figure 2A). This is likely driven by specific subtypes of colonocytes, as we found that *CA1*+ colonocytes (Figure 2B), especially *CA1*+ late colonocytes (Figure 2C), are significantly increased in abundance in CD relative to NIBD. We observed only a slight difference in the relative abundance of mature absorptive cells (combined *CA1*+ early, *CA1*+ late, and *CEACAM7*+ colonocytes) vs. mature secretory cells (combined mature goblet cells and EECs) (Figure 2D). However, between the two major secretory lineages, we discovered a significant skew toward the mature goblet fate in CD (Figure 2E). Even within the goblet lineage we observed a shift toward heightened goblet maturation in CD compared to NIBD (Figure 2F).

**Figure 2:**
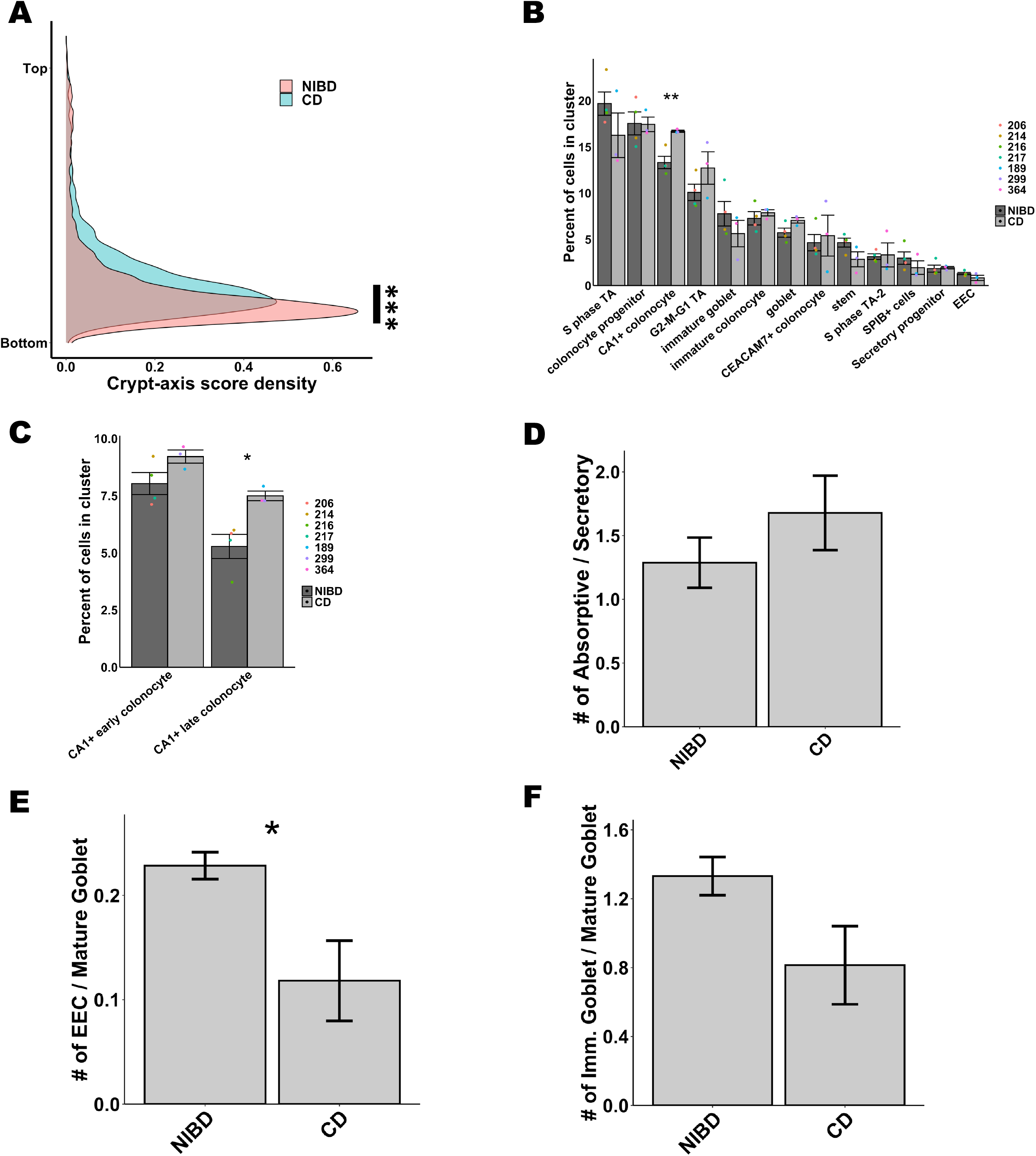
Crohn’s disease causes shifts in colonic epithelial landscape. (A) Crypt-axis score density for NIBD and CD cells separately reveals a shift toward the colonic crypt top in CD. Significance of shift was determined using Kolmogorov-Smirnov test (p < 0.001 = ***) (B) Mean cell abundances across NIBD and CD samples reveals significant increase in CA1+ colonocytes in CD. Dots show abundance values of individual samples. (C) Mean cell abundances for either CA1+ early or late colonocyte clusters across NIBD and CD samples reveals a significant increase only in the CA1+ late colonocytes. Dots show abundances of individual samples. Abundances of other clusters not shown. (D) Mean ratio of absorptive to secretory cells across NIBD and CD samples. (E) Mean ratio of EEC to mature goblet cells across NIBD and CD samples reveals a significant shift in the secretory lineage toward mature goblet cells in CD. (F) Mean ratio of immature goblet to mature goblet across NIBD and CD samples reveal a shift toward mature goblet cells in CD. If not indicated, p-values are calculated by a Student’s t-test (p < 0.05 = *, p < 0.01 = **). NIBD = Non-inflammatory bowel disease, CD = Crohn’s disease, EEC = enteroendocrine cells.

### Different colonocyte clusters exhibit unique changes in gene expression in Crohn’s disease

To investigate the genes and pathways most altered in CD in a cell-type specific manner, we first performed differential gene expression analysis in each of the 14 cell types separately. We identified a varying number of significantly (adjusted P < 0.05) differentially expressed genes (DEGs) across cell types (Figure 3A). We found that the number of DEGs are roughly proportional to the number of cells in each of the clusters (Figure S2A), indicating that the variation in number of DEGs is due at least in part to variability in statistical power. A notable exception is the *CA1*+ late colonocyte cluster, which exhibits as many or more DEGs than several clusters with a greater number of cells, including the *CA1*+ early colonocyte, G2-M-G1 TA, immature and mature goblet, and immature colonocyte clusters. Also, even though there are >200 fewer cells in the stem cell cluster compared to the mature goblet cell cluster, the stem cells exhibit more DEGs than mature goblet cells (Figure 3A), pointing to a robust change in the molecular profile of stem cells in CD. To investigate cell type specific changes in gene expression in the absorptive and secretory lineages, we first focused on two major colonocyte clusters (*CA1*+ late colonocyte and *CEACAM7*+ colonocyte) and two secretory clusters (mature goblet cells and EECs).

**Figure 3:**
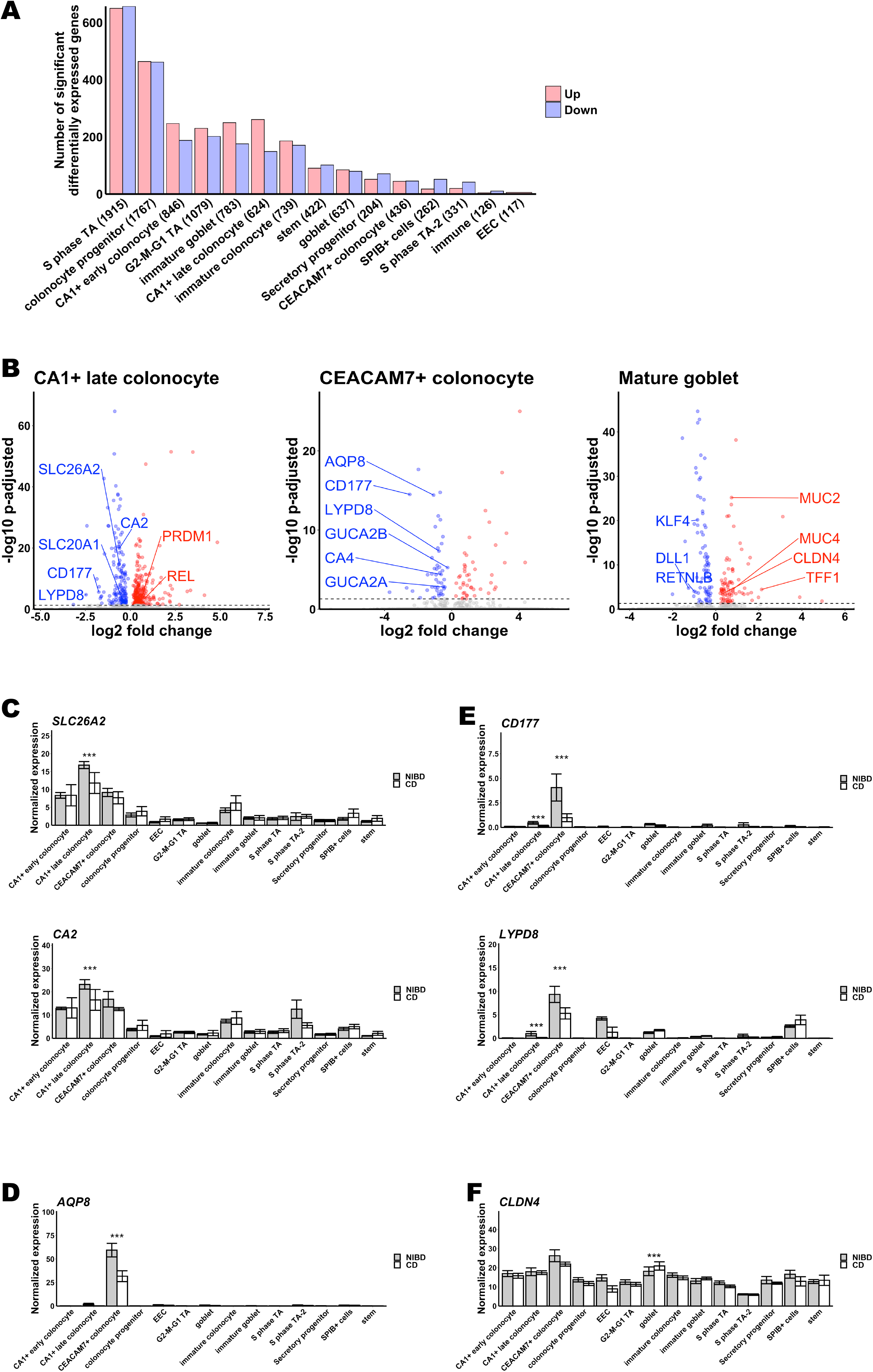
Effect of Crohn’s disease within cell types. (A) Number of significant differentially expressed genes in CD versus NIBD. Clusters are shown on the x-axis followed by the number of cells within the cluster in parenthetical notation and the number of differentially expressed genes is shown on the y-axis. (B) Differentially expressed genes for three clusters; CA1+ late colonocyte, CEACAM7+ colonocyte, and mature goblet. Log2 fold change is shown on x-axis and adjusted p-value is shown on the y-axis. (C-F) Mean expression of *SLC26A2* & *CA2* (C), *AQP8* (D), *CD177 & LYPD8* (E), and *CLDN4* (F) across clusters for NIBD and CD samples demonstrates the specificity of expression and differential expression within clusters. P-values are calculated by Wilcoxon Rank Sum test (adjusted p < 0.05 = *, adjusted p < 0.01 = **, adjusted p < 0.001 = ***). NIBD = Non-inflammatory bowel disease, CD = Crohn’s disease.

We found that 45 genes (30 up, 15 down in CD relative to NIBD) are significantly altered uniquely in *CA1*+ late colonocytes (Table S2). Among those significantly up-regulated uniquely in *CA1*+ late colonocytes are *PRDM1* and *REL*, and among those down-regulated uniquely are *CA2, SLC26A2*, and *SLC20A1* (Figure 3B). *PRDM1* and *REL* have been implicated in anti-inflammatory and microbial-sensing pathways in the colon, and have been reported as harboring variants associated with CD in genome-wide association studies (GWAS) (Ellinghaus et al., 2013; Khor et al., 2011); however, the increased expression in CD in only this subtype of colonocytes has not been demonstrated previously. *CA2* and *SLC26A2* are not only uniquely down-regulated in *CA1*+ late colonocytes, but are also most highly expressed in this cluster (Figure 3C), suggesting that normal functions of this cell type such as anion transport are compromised in CD (SLC26A2 and SLC20A1 are sulfate and phosphate transporters, respectively, contributing to solute homeostasis in the colon).

There are 29 genes (7 up, 22 down) significantly altered uniquely in *CEACAM7*+ colonocytes (Table S3). Among those down-regulated are *CA4, AQP8, GUCA2A*, and *GUCA2B* (Figure 3B), each of which has been implicated in various normal functions of the colonic epithelium, such as the role of AQP8 in colonic epithelial water transport (Masyuk et al., 2002). Decreased colonic epithelial expression of *AQP8* has been reported in IBD previously (Ricanek et al., 2015), and has been suggested as a candidate therapeutic target for diarrheal diseases (Escudero-Hernandez et al., 2020). Here, we show that the gene that codes for this important IBD-related protein is dramatically down-regulated in CD in *CEACAM7*+ colonocytes only and not in any other cell type of the colonic epithelium (Figure 3D).

We found that 4 genes (1 up, 3 down in CD relative to NIBD) are significantly altered in both *CA1*+ colonocytes and *CEACAM7*+ colonocytes, and in no other cell type. Shared down-regulated genes include *CD177* and *LYPD8* (Figure 3B), both of which are in the same family of proteins containing the LY6/PLAUR domain. The latter encodes a protein that protects the gut from microbial invasion and is critical for maintaining barrier integrity and preventing intestinal inflammation (Hsu et al., 2017; Okumura et al., 2020; Okumura et al., 2016). *LYPD8* is expressed 5-fold greater in *CEACAM7*+ colonocytes compared to *CA1*+ late colonocytes, but it is significantly down-regulated in CD in both cell types (Figure 3E). Surprisingly, it is also highly expressed and dramatically reduced in CD in EECs (Figure 3E), though this decrease doesn’t achieve significance likely due to the very small number of EECs, contributing to low statistical power.

### The mature goblet program is enhanced whereas enteroendocrine drivers and L-cell markers are suppressed in Crohn’s disease

Goblet cells secrete mucins to create a protective barrier for the colon from luminal content. In mature goblet cells, classic gene markers of maturity and function, including *MUC2, MUC4*, and *TFF1* are significantly elevated in CD, whereas markers of immaturity, including *KLF4, DLL1*, and *RETNLB*, are significantly reduced in CD (Figure 3B). This is concordant with our finding of a shift toward heightened goblet maturation in CD compared to NIBD (Figure 2E). Interestingly, levels of *CLDN4*, previously reported to be highly expressed in EECs (Nagatake et al., 2014)and also studied in the context of colonocyte barrier function (Watari et al., 2017), are significantly elevated in the mature goblet cell cluster while modestly reduced in both *CEACAM7*+ colonocytes and EECs (Figure 3F). In fact, in CD, the levels of *CLDN4* in mature goblet cells rise to what is observed in *CEACAM7*+ colonocytes (Figure 3F), the cells in which *CLDN4* is most highly expressed in NIBD.

EECs, which secrete hormones in response to nutrients in order to maintain metabolic homeostasis, are not well-studied in CD. To examine the change in the molecular character of EECs in CD, we first examined the genes encoding key transcription factors (n=10) that contribute to EEC maturation. We found that seven out of the ten are significantly altered in CD, all of which are down-regulated (Figure S3A), which is consistent with our observation that the abundance of EECs (relative to mature goblet) trends lower in CD compared to NIBD (Figure 2E). We next evaluated the genes encoding the major hormones (n=7) that are produced and secreted from EECs. We found that two in particular, *GCG* and *PYY*, both of which are expressed in the colonic L-cell subtype of EECs (Billing et al., 2018), are more than 3-fold reduced in CD vs. NIBD (Figure S3B). *GCG* encodes the key metabolic hormone GLP-1 that promotes systemic energy homeostasis (Barrera et al., 2011), and *PYY* encodes a signal that promotes satiety. Notably, the gene *Nts*, which encodes the pro-inflammatory peptide neurotensin (Castagliuolo et al., 1999), is highly up-regulated in CD (Figure S3B).

### The canonical colonic stem cell signature is disrupted in Crohn’s disease

Dysfunction in the intestinal stem cell population (ISC) in Crohn’s disease has been proposed but not rigorously evaluated and documented. To investigate this possibility, we analyzed DEGs in the ISC population in CD relative to NIBD. We found that in ISCs, among the most highly up-regulated genes in CD are *PLA2G2A* and *KLF6* (Figure 4A), both of which encode factors that are known to negatively regulate Wnt signaling in the crypts (Cheung et al., 2020; Schewe et al., 2016). Although we found that *PLA2G2A* is up-regulated in many other clusters also, *KLF6* is significantly elevated primarily in ISCs. Accordingly, we observed that numerous genes in the Wnt signaling pathway (including *CDCA7, CDK6, CCDC115, MYC, RNMT, TGIF1, YBX1*, and *FOS*) are significantly reduced in expression in the ISC cluster in the CD samples relative to NIBD (Figure 4B). Moreover, in the case of *CCDC115* and *RNMT*, they are significantly altered only in ISCs and not in any other cluster (Figure 4C).

**Figure 4:**
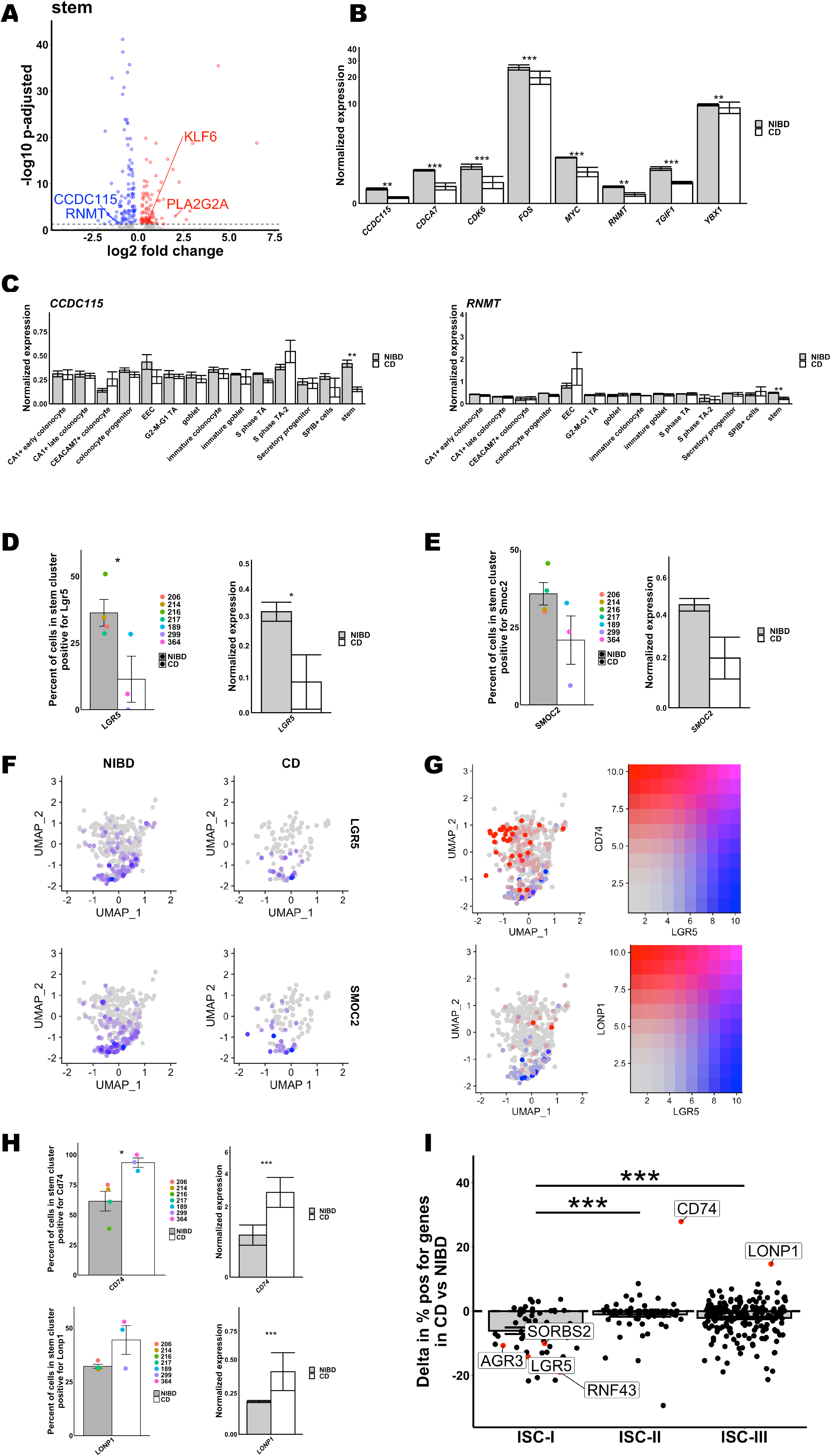
Crohn’s disease disrupts stem cell homeostasis. (A) Differentially expressed gene within the stem cluster. Log2 fold change is shown on x-axis and adjusted p-value on the y-axis. (B) Mean expression of eight mediators of the Wnt pathway in NIBD and CD samples reveal a consistent decrease in response to CD. (C) Mean expression of *CCDC115 & RNMT* across clusters shows a significant decrease only occurs in the stem cluster. (D-E) Percent of cells positive for *LGR5* (D, left) and *SMOC2* (E, left) or the average normalized expression for *LGR5* (D, right) and *SMOC2* (E, right) in stem cluster for NIBD or CD samples. Dots relate to individual sample value. (F) UMAP of the stem cluster with expression of *LGR5* or *SMOC2* overlain, separated by NIBD/CD. (G) UMAP of stem cluster with *LGR5* expression (red) co-overlain with either *CD74* (top) or *LONP1* (bottom) (both blue) expression. Scale for expression is shown on right. (H) Percent of cells positive for *CD74* (left top) and *LONP1* (left bottom) or the average normalized expression for *CD74* (right top) and *LONP1* (right bottom) in stem cluster for NIBD or CD samples. Dots relate to individual sample value. (I) Shift in percent positive of genes in either ISC-I, ISC-II, or ISC-III cell types between NIBD and CD. P-values for expression bar plots was calculated using the Wilcoxon Rank Sum test and the p-value for the ISC subtype shifts was calculated using a Student’s t-test (p < 0.05 = *, p < 0.01 = **, p < 0.001 = ***). NIBD = Non-inflammatory bowel disease, CD = Crohn’s disease, ISC = intestinal stem cell, UMAP = Uniform Manifold Approximation and Projection.

We next assessed whether the Wnt-responsive, canonical marker of colonic stem cells, *LGR5* (Barker et al., 2007), is affected in CD. We found that both the percentage of *LGR5*+ cells and the expression of *LGR5* in the ISC cluster are significantly reduced in CD relative to NIBD (Figure 4D). As a confirmation of this finding, we also detected a similar trend for *SMOC2* (Figure 4E), which is known to be specifically enriched in the *LGR5*+ stem cell compartment (Munoz et al., 2012).

Intriguingly, we noticed that not all of the cells in the ISC cluster exhibit high expression of *LGR5* or *SMOC2*, even in the NIBD samples (Figure 4F). Previous work has distinguished three subtypes of ISCs: ISC-I, ISC-II, and ISC-III. ISC-I comprises canonical *LGR5*-high stem cells, whereas ISC-II and ISC-III are composed of *LGR5*-low stem cells that are more proliferative, more differentiated, and potentially antigen-presenting (Biton et al., 2018). We analyzed established markers (Biton et al., 2018) of the ISC-II and ISC-III subtypes, *CD74* and *LONP1*, respectively, and confirmed that they are indeed present almost exclusively in the cells of the ISC cluster that are *LGR5*-low or *LGR5*-negative (Figure 4G). Moreover, we observed that both the percentage of *CD74*+ cells and the expression of *CD74* in the ISC cluster are significantly increased in CD relative to NIBD (Figure 4H). A similar trend was observed for the ISC-II marker gene *LONP1* (Figure 4G,H). We also found that the vast majority of ISC-I marker genes (including but not limited to *LGR5*) are reduced in representation in CD ISCs (Figure 4I). Taken together, these data show that there is a shift away from an ISC-I signature in CD, indicative of a disrupted, non-homeostatic state in the colonic crypts.

### Detailed analysis of the SPIB+ cell cluster reveals new, rare cell types altered in Crohn’s disease

The least understood cell type that we have identified is the *SPIB*+ cluster. To dissect this further, we performed sub-clustering within this cluster only, and identified 4 sub-clusters (Figure 5A). We found that each of these sub-clusters expresses a unique set of markers (Methods, Figure 5B,C). Notably, the gene *OTOP2* specifically marks *SPIB*+ sub-cluster 1; *BEST4* (associated specifically with absorptive cells (Ito et al., 2013)) and *CA7* are present only in *SPIB*+ sub-clusters 0 and 1; *SLC12A2* is enriched in *SPIB*+ sub-clusters 2 and 3; and *LYZ* is elevated in *SPIB*+ sub-cluster 3, though it is not highly specific (Figure 5B,C). The crypt-axis score analysis revealed a locational gradient of SPIB+ sub-clusters (Figure 5D). Only one of the sub-clusters (sub-cluster 1) is closer to the crypt-top, whereas the others are closer to crypt-bottom (Figure 5D). Notably, we found that the relative abundance of *SPIB*+ sub-cluster 1 (*OTOP2*+/*CA7*+) is significantly elevated in CD as compared to NIBD (Figure 5E).

**Figure 5:**
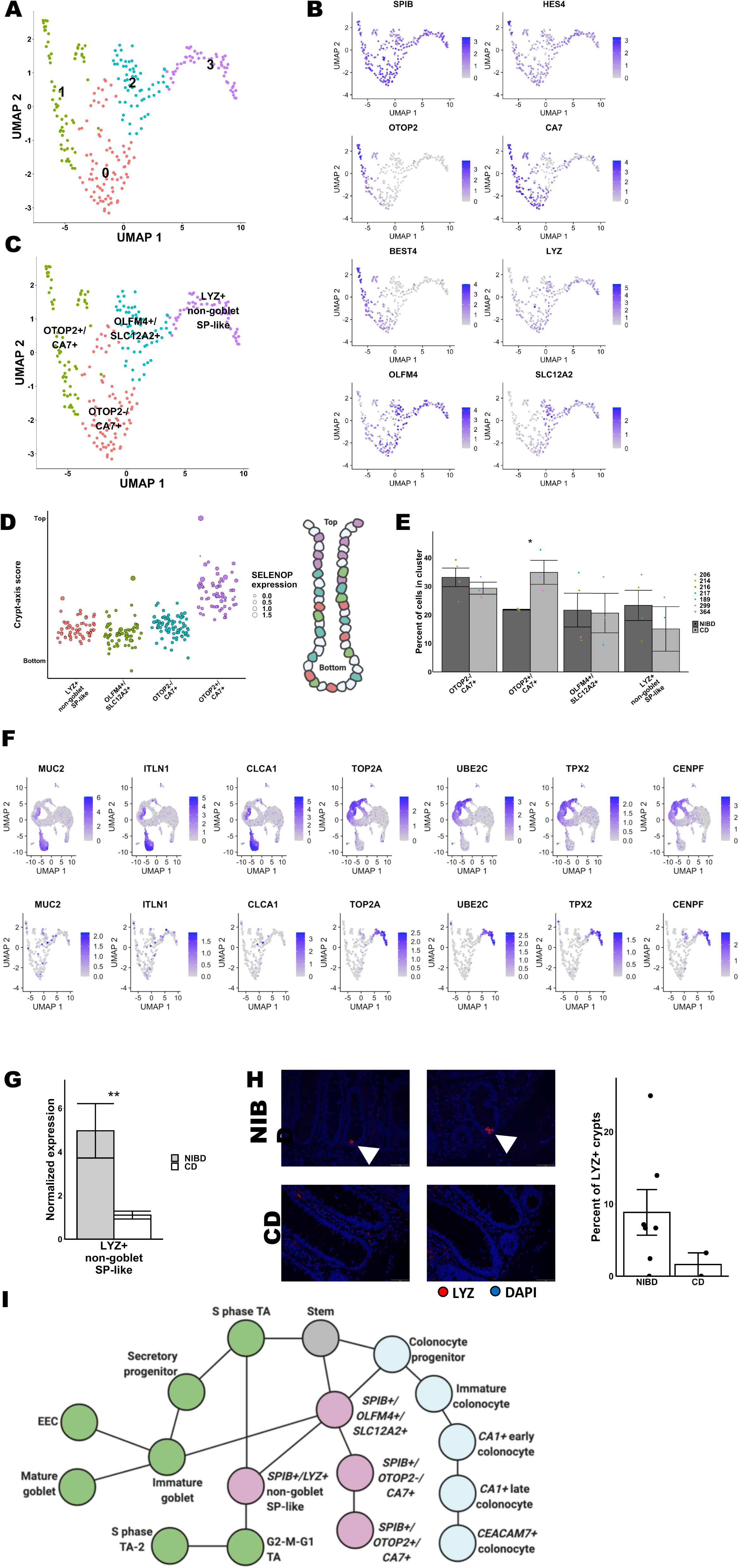
Subcluster analysis of SPIB+ cells reveals distinct lineages. (A) UMAP following subclustering of SPIB+ cells with labeling of the four identified subclusters. (B) UMAP of SPIB+ cell subclusters overlain with gene expression from genes marking the entire cluster or distinct subclusters. (C) SPIB+ cell subcluster UMAP following cell type assignment using highly enriched markers of the subclusters. (D) Crypt-axis scores (low near crypt bottom, high near crypt top) of subcluster-assigned cells uncovers a colonocyte signature in the OTOP2+/CA7+ subcluster. Clusters are arranged on x-axis by mean crypt-axis score. The size of the dot corresponds to level of expression of *SELENOP*, a known marker of the top of the crypt. (E) Mean cluster cell abundances across NIBD and CD samples reveals significant increase in OTOP2+/CA7+ subcluster in CD. Dots show abundances of individual samples. (F) UMAP of either all clusters (top) or SPIB+ cell subclusters (bottom) overlain with the expression of markers of secretory progenitor cells. (G) Mean normalized expression of *LYZ* in NIBD and CD samples in the LYZ+ non-goblet SP-like subcluster. (H) LYZ-detected immunofluorescence in colonic crypts from either NIBD or CD patients examples (left) and quantified by sample (right). Mean percent of crypts positive for LYZ in NIBD or CD. (I) Lineage reconstruction as determined by PAGA. NIBD = Non-inflammatory bowel disease, CD = Crohn’s disease, UMAP = Uniform Manifold Approximation and Projection, PAGA = partition-based graph abstraction.

We also noted that several markers of proliferation, particularly those that also mark the secretory progenitor (SP) cells, are enriched in *SPIB*+ sub-cluster 3 relative to the other *SPIB*+ sub-clusters. Among the 7 most specific markers of SP cells (Methods), we found that 4 (*TOP2A, UBE2C, TPX2*, and *CENPF*) are dramatically enriched in *SPIB*+ sub-cluster 3 relative to the other *SPIB*+ sub-clusters (Figure 5F). The remaining three, *CLCA1*, MUC2, and ITLN1, which are not found in *SPIB*+ sub-cluster 3, also mark immature and mature goblet cells (Figure 5F). Therefore, we labeled *SPIB*+ sub-cluster 3 as *LYZ*+ non-goblet SP-like cells (Figure 5D). Notably, *LYZ* expression in these cells is significantly reduced in CD relative to NIBD (Figure 5G). We validated the presence of LYZ in crypt-bottom cells in NIBD samples and the depletion of this signal in CD samples by immunohistochemistry (Figure 5H). The lineage relationship among the SPIB+ sub-clusters and other cluster is shown in Figure 5I.

### Genes implicated in Crohn’s disease risk exhibit cell-type specific patterns of aberrant expression within the colonic epithelium

To investigate the colonic epithelial expression of genes implicated in CD, we first defined a list of genes (n=261) nearest to every genetic variant significantly associated with CD based on GWAS, of which 208 were detected in this dataset (Methods). For each of these genes we determined the percentage of cells positive and average expression across cells within each of the 14 colonic epithelial cell clusters (Figure 6A). We confirmed that genes expected to be enriched in the lamina propria, such as *IL10* and *NOD2*, are not detected robustly in any of the epithelial cell clusters (Figure 6A). Indeed, several other genes fell into this category as well, including *IL2RA, LTA, TNFSF8, CD244*, and *NELL1*, suggesting that if these genes are involved in CD pathophysiology, their primary roles are most likely outside of the epithelium. We observed that some CD genes are detected at comparable levels across many epithelial cell clusters, such as *ATG5, SKAP2, TAGLN2, PPM1G*, and *PRDX5*, whereas other genes are enriched in only one or few cell types, such as *ITLN1* (mature goblet cells) (Figure S4), *CACNA2D1* (EECs), *COL5A1* (EECs), *RIPOR1* (EECs), *IFNGR2* (*CEACAM7*+ colonocytes), *NOTCH2* (*SPIB*+ cells), and *ATG16L2* (*SPIB*+ cells). Investigating the latter two further within the *SPIB*+ cluster, we found that while *NOTCH2* is ubiquitously expressed across all *SPIB*+ sub-clusters, *ATG16L2* is enriched in *SPIB*+ sub-cluster 1 (*LYZ*+ non-goblet SP-like cells) (Figure 6B).

**Figure 6:**
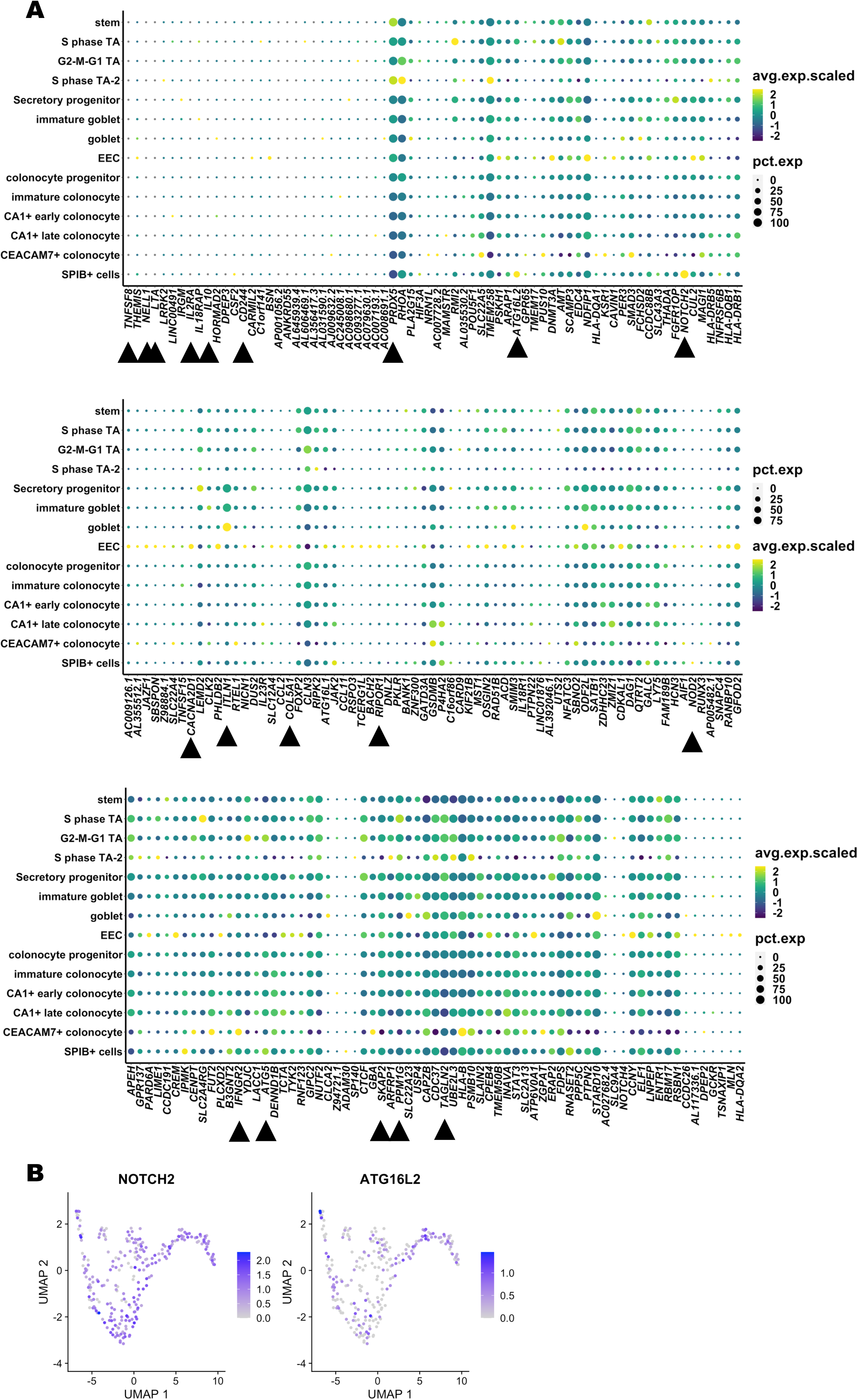
Expression of CD-associated risk genes across clusters. (A) Dot plots showing expression and percentage of cells expressing CD-associated risk genes (columns) across clusters (rows). For each gene, color of dot corresponds to the average scaled expression and the size of the dot corresponds to the percent of cells in cluster expressing the gene. (B) UMAP of SPIB+ cells subclusters overlain with gene expression from *NOTCH2* or *ATG16L2*. NIBD = Non-inflammatory bowel disease, CD = Crohn’s disease, UMAP = Uniform Manifold Approximation and Projection.

We next sought to determine whether any CD-associated genes are significantly altered in expression in CD relative to NIBD in one or more epithelial cell clusters. Some genes exhibit significant changes in gene expression across many clusters (Figure 7A), including HLA family genes such as *HLA-DQB1, HLA-DRB1*, and *HLA-DRB5*. These data point to the systematic increase in CD genes encoding for major histocompatibility complex (MHC) class II factors, generally active in antigen-presenting cells, across most epithelial cells including stem cells as described previously (Biton et al., 2018), with some minor exceptions (e.g., EECs for *HLA-DRB5* or *CA1*+ early colonocyte for *HLA-DQB1*). Most genes that change in CD exhibit cell-type specific patterns. Notable examples include genes that are nearly specifically altered in EECs, including *CACNA2D1, COL5A1, RIPOR1, RNF123*, and *LEMD2*. The first three of these are more highly expressed in EECs in NIBD relative to other cell types (Figure 6A), whereas *RNF123* and *LEMD2* expression levels are comparable across most cell types in NIBD but uniquely suppressed in EECs in CD patients (Figure 6A,7A).

**Figure 7:**
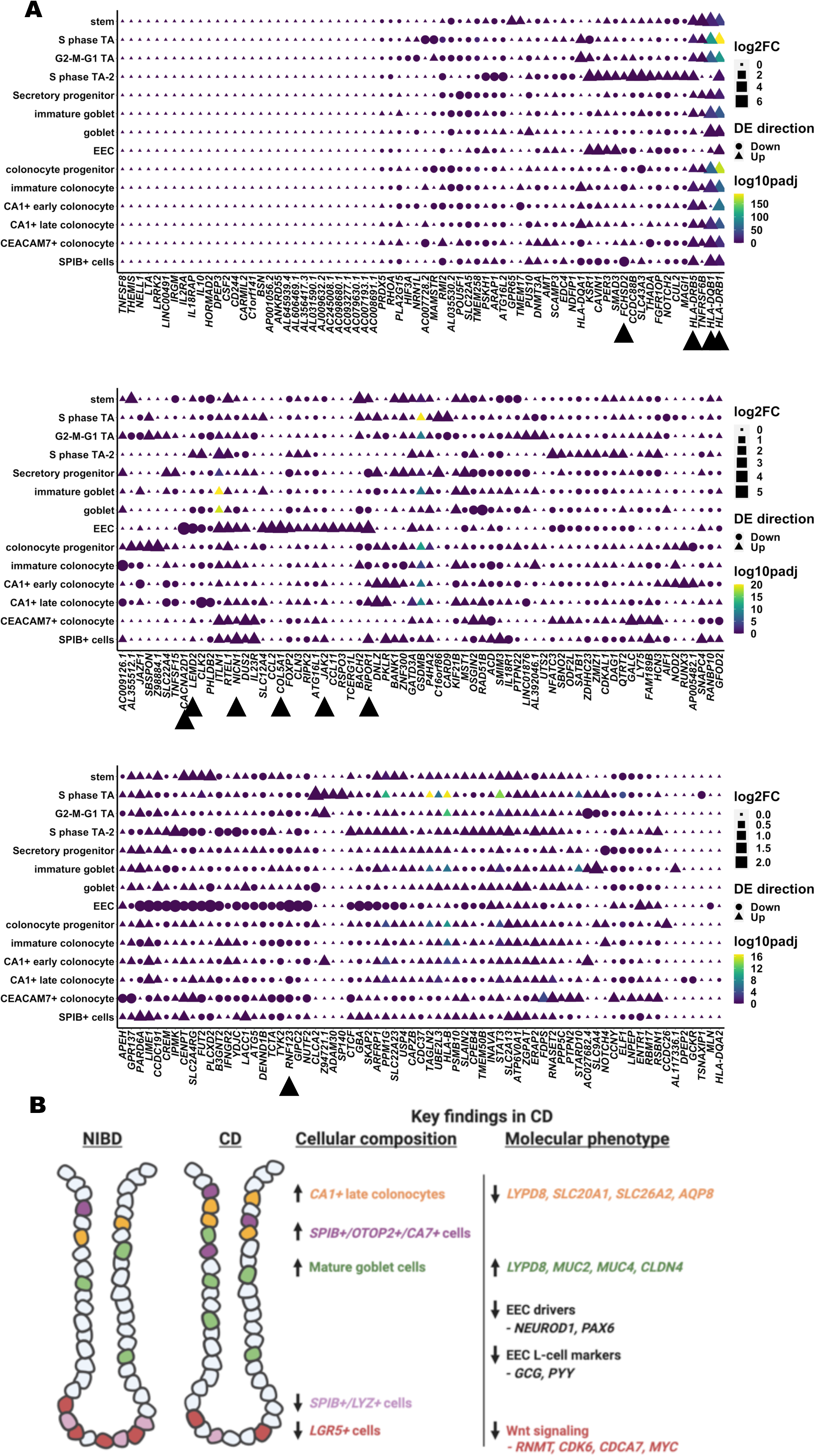
Differential expression of CD-associated risk genes across clusters. (A) Dot plots showing the log2 fold change of CD compared to NIBD within clusters, the direction of change, and the statistical significance of CD-associated risk genes (columns) across clusters (rows). For each gene, color of the dot corresponds to the log2 fold change, the shape of the dot corresponds to the direction of change, and the color of the dot corresponds to the adjusted p-value of the comparison. Adjusted p-value was calculated from the Wilcoxon Rank Sum test. NIBD = Non-inflammatory bowel disease, CD = Crohn’s disease.

Other examples include genes preferentially altered in stem cells, including *BACH2, GPR65*, and *TNFRSF6B*; in mature goblet cells, most notably *ITLN1*; as well as in *SPIB*+ cells, including *FCHSD2* and *NICN1*. In the case of *FCHSD2*, it is suppressed across all *SPIB*+ sub-clusters, whereas the change in *NICN1* is driven almost exclusively by *SPIB*+ sub-cluster 2. We found that the CD gene *JAK2* is also significantly elevated in *SPIB*+ cells (specifically, sub-cluster 3), though this aberration is shared with EECs as well (Figure 7A). The functions of *FCHSD2* and *NICN1* in CD pathophysiology are currently unknown, whereas *JAK2* is well-studied in CD risk and JAK inhibitors are currently FDA-approved for IBD. Notably, though, the connection being made here between *JAK2* and EECs or *SPIB*+ cells in the context of CD has not been reported previously.

## Discussion

To our knowledge, this study represents the first single-cell analysis of colonic epithelial cells from treatment-naïve adult CD patients. A unique feature of this study is that the samples were isolated from treatment-naïve individuals, which means that the results are not confounded by the effects of drugs. Importantly, we specifically avoided macroscopically inflamed tissue in order to focus on cellular reprogramming that occurs more generally in the epithelial layer within the colon during CD. Inflammation can also lead to destruction of the epithelial layer, making it an unreliable indicator of IEC gene signatures. We identified almost 15 different types of epithelial cells, including several different subtypes of colonocytes and colonocyte progenitors. Each of the major clusters that express mature colonocyte markers, *CEACAM7*+ cells, *CA1*+ late colonocytes, and *SPIB*+ cells, express a unique set of genes. Notably, we found that *CEACAM7*+ cells are marked by *LINC01133*, which as we understand is the first reported lncRNA marker of a colonic epithelial cell type. Although *LINC01133* has been implicated in the control of tumor phenotypes in colon cancer (Kong et al., 2016), its role in the normal function of *CEACAM7*+ colonocytes remains unknown and merits further investigation. Also, among the colonocyte clusters, the *CA1*+ late cluster is the only colonocyte subtype for which relative abundance is significantly altered in CD, and this too warrants study in the future.

Study of cellular composition revealed an increase in late-stage colonocytes in CD compared to healthy controls. However, in-depth analysis of the single cell data revealed aberrancies in the molecular character of these cells. For example, we observed dramatic reduction of solute and water transporters, suggestive of compromised colonocyte function in CD. Particularly notable is the dramatic decrease in *AQP8* only in *CEACAM7*+ colonocytes. Although *AQP8* has previously been shown to be suppressed in IBD (Ricanek et al., 2015), our study demonstrates for the first time that this effect is very likely driven by one specific colonocyte subtype. Furthermore, the significant reduction in CD of *LYPD8*, which encodes an important anti-microbial factor (Hsu et al., 2017; Okumura et al., 2020; Okumura et al., 2016), in both *CEACAM7*+ colonocytes and *CA1*+ late colonocytes, is a possible indication of the beginning stages of impaired colonocyte contribution to barrier function. We also show that *LYPD8* is reduced in expression in EECs from CD patients, which has not been demonstrated previously.

Of note, we find that *LYPD8* is modestly, albeit not significantly, increased in the mature goblet cell cluster. Also, mature goblet cells show both increased abundance in CD as well as increased expression of genes that code for mucins, including *MUC2* and *MUC4*. This result is consistent with another recent study (Elmentaite et al., 2020), which reported a significant increase in goblet cells in pediatric ileal CD samples. This finding, coupled with alterations in the molecular phenotype of mature colonocytes, raises the possibility of a compensatory response of mature goblet cells to the potentially weakened function of colonocytes in CD. This notion is further supported by significantly increased *CLDN4* expression in mature goblet cells in CD, which encodes for a tight-junction protein that promotes barrier integrity (Watari et al., 2017), to levels that match what is observed in *CEACAM7*+ colonocytes (where *CLDN4* expression is normally most prominent).

We found a novel shift in ISCs away from the canonical *LGR5*+ signature in CD compared to NIBD, which motivates new avenues of future investigations into the dysfunction in ISCs as well as in lineage determination and colonic epithelial renewal during CD development. Among the numerous pro-Wnt signaling marker genes with reduced expression in ISCs in CD, two are uniquely suppressed only in ISCs, *CCDC115* and *RNMT*. RNMT is recruited to Wnt signaling gene promoters by MYC (Posternak et al., 2017), which we show is suppressed in ISCs in CD, and therefore may be involved in mediating the shift away *LGR5*+ ISCs. To our knowledge, *CCDC115* has not been studied previously in ISCs and may represent a novel factor in their function and CD etiology. We also found increased expression of MHC class II factors such as *CD74, HLA-DQB1*, and *HLA-DRB1*, normally associated with antigen presentation, in ISCs of CD patients. A recent study showed that murine small intestinal stem cells act as non-conventional antigen-presenting cells to activate lamina propria T cells especially under certain conditions such as enteric infection (Biton et al., 2018). Whether this also occurs in CD is not known and merits further investigation.

One of the most intriguing findings in our study is that an initial cluster characterized by near ubiquitous *SPIB, HES4*, and *NOTCH2* expression is more accurately annotated as four distinct sub-clusters based on unique markers that clearly distinguish each. Among these sub-clusters, the *OTOP2*+/*CA7*+ sub-cluster, is significantly increased in relative abundance in CD compared to NIBD. Two previous single cell studies (Parikh et al., 2019; Smillie et al., 2019) of ulcerative colitis (UC) also identified this cell type but actually found that its abundance is decreased in UC patients compared to healthy controls. There are several possible explanations for this difference in our study: CD has a different etiology than UC; our analysis is focused on macroscopically non-inflamed regions; and our treatment-naïve patients may be in a different stage of disease course. These possibilities point to the importance of future single cell studies that carefully investigate differences between CD vs. UC and also include longitudinal assessments along disease course. We also identified a sub-cluster of *SPIB*+ cells, comprising *LYZ*+ cells. Notably, *LYZ* expression is reduced and *LYZ*+ cells are depleted in CD compared to NIBD, both at the RNA and protein level. The *SPIB*+/*LYZ*+ cells may correspond to previously annotated Paneth-like cells (PLCs)(Wang et al., 2020); however, besides *LYZ*, we do not detect any classic markers of Paneth cells and therefore we termed them “non-goblet secretory progenitor-like cells” based on their gene signature.

Finally, in this study, we link CD GWAS risk genes to specific epithelial cell subtypes in which they are detected and/or aberrantly expressed. These discoveries may offer clues about the potential molecular mechanisms by which these genes contribute to CD etiology. For example, *ATG16L2* is enriched in the *SPIB*+ cluster, particularly within the *SPIB*+/*LYZ*+ sub-cluster, suggesting that this gene may contribute to CD etiology through the novel functions of this uncharacterized cluster, which may center on anti-microbial functions given the presence of *LYZ*. We also found that increased *JAK2* expression is prominent not only in the *SPIB*+ cluster but also in EECs, which is completely unexpected and merits more detailed functional investigation given that JAK inhibitors are approved for use in ulcerative colitis and are in clinical trials for CD. Overall, we believe that this study offers a unique picture, at unprecedented resolution, of the cellular and molecular landscape of the colonic epithelium in treatment-naïve adult CD.

Future single-cell studies must expand on this work to include larger cohorts and to assess the impact of age, gender, CD sub-type (Keith et al., 2018; Weiser et al., 2018), disease region (e.g., ileum vs. colon), disease duration, and treatment history on cell composition and molecular phenotype. Longitudinal investigations to uncover changes to the cellular landscape and molecular phenotype over the course of disease progression are also merited. Finally, it will be exciting in the future to explore the relationship between specific luminal bacterial changes and alterations at the single-cell level, given the intimate relationship between microbial dysbiosis and CD pathogenesis.

## Methods

### Ethical statement

The study was conducted in accordance with the Declaration of Helsinki and Good Clinical Practice. The study protocol was approved by the institutional review board at University of North Carolina at Chapel Hill (approval number: 19-0819 and 17-0236). All participants provided written informed consent prior to inclusion in the study. All participants are identified by number and not by name or any protected health information.

### Single cell RNA-sequencing

Colonic mucosa was obtained endoscopically as biopsies from patients with treatment-naive CD and NIBD healthy controls. Cross-sectional clinical data was collected at the time of sampling. All samples were collected from regions of ascending colon without macroscopic inflammation. Isolation of colonic primary IECs was performed as reported previously (Toyonaga et al., 2020; Wang et al., 2017). This method has previously been shown to result in >95% purity of IECs (Camp et al., 2014). Single cell libraries were constructed using the Chromium Single Cell 3’Reagent Kits (V3) according to manufacturer instructions.

### Single cell transcriptome data analysis

Sample single cell fastqs were aligned to the human genome (GRCh38-3.1.0) using 10x Genomics cellranger count (v4.0.0) to obtain gene/cell count matrices. Sample filtering, normalization, integration, clustering and visualization was accomplished using Seurat (v3.2). To maintain sample quality, cells with less than 1000 detected genes or greater than 25% of reads aligning to mitochondrial genes were removed. Additionally, cells with greater than 50000 reads were removed as potential doublets. To control for read depth, sample counts were log normalized. Samples were merged using the Seurat integration anchor workflow based on the 2000 most variable genes. Clustering and identification of nearest neighbors relied on 11 PCA dimensions. Cells were clustered at a resolution of 0.7. To focus on epithelial cells, immune clusters were identified by expression of known immune cell type markers. Immune clusters were removed, and the integration workflow was repeated using the 2000 most variable genes of the remaining epithelial cells. Clustering and identification of nearest neighbors relied on 10 PCA dimensions. Epithelial cells were clustered using a resolution of 0.8. Highly enriched markers for each cluster were determined using the Seurat function FindAllMarkers, which compares the gene expression within a cluster to all other clusters. Genes that are up-regulated with a log fold change greater than 0.25 and a Wilcoxon Rank Sum test p-value < 0.01 were considered highly enriched markers. Clusters were assigned cell types based on expression of known markers and/or highly enriched markers of the cluster. Despite earlier filtering, one of the resulting clusters was immune cells and therefore removed from the analysis. Samples were found to have similar count, genes, and mitochondrial read percentage per cell (Table S4). Differential expression of genes between CD and NIBD samples was determined within clusters using the Wilcoxon Rank Sum test.

Sub-clustering of SPIB+ cells was performed to identify discrete cell types within the cluster. The SPIB+ cells were sub-clustered using the integrated counts. Clustering and identification of nearest neighbors relied on 10 PCA dimensions. SPIB+ cells were sub-clustered using a resolution of 0.8. Assignment of cell types to sub-clusters was determined by the highly enriched markers of the subcluster.

### Determination of highly enriched marker genes

Highly enriched genes were determined by one of two different thresholds. In both thresholds, the genes had to be up-regulated >0.5 log fold change in the cluster of interest vs. all other clusters. Using the stringent threshold, a gene had to be expressed in greater than 90% of the cells in the cluster of interest and less than 30% in all other clusters (Figure 5F). Using the slightly relaxed threshold, a gene had to be expressed in greater than 80% of cells in the cluster of interest and less than 40% of cells in all other clusters (Figure 1D).

### Crypt-axis score

The crypt-axis score was assigned for each cell and was based on the expression of a previously defined set of genes: *SELENOP, CEACAM7, PLAC8, CEACAM1, TSPAN1, CEACAM5, CEACAM6, IFI27, DHRS9, KRT20, RHOC, CD177, PKIB, HPGD, LYPD8* (Parikh et al., 2019). For each gene, expression within a cell was divided by the max expression across all cells to mitigate the weight of highly expressed genes. The max expression normalized values places the gene’s expression on a 0 to 1 scale. The crypt-axis score is the summation across all genes of the max-normalized expression values.

### Intestinal stem cell analysis

Genes constituting the three classes of intestinal stem cells (ISC) were obtained from Biton et al (Biton et al., 2018). In the stem cluster, the percent of NIBD or CD cells positive for the genes was calculated and the delta was determined by subtracting the NIBD percent positive from the CD percent positive for each gene. The overall shift in an ISC class was determined by averaging the delta across all genes within the ISC class.

### Integrative analysis with IBD genome-wide association study results

A list of 5743 Crohn’s disease risk-associated SNPs were identified through GWAS. The closest genes to the disease risk-associated SNPs were identified using bedtools closest (v2.27), resulting in 261 unique genes. Of these genes, 208 were present in the filtered dataset. For the figures, the gene order was determined by hierarchical clustering of the Euclidean distance of the log2 fold change for each gene across clusters.

### Velocyto

The velocity pseudotime figure was generated using scVelo (v0.2.2). The velocities were computed using the dynamical model, and we performed a likelihood-ratio test to test for differential kinetics between clusters and corrected the velocity for differential kinetics. The root index was set as the first indexed stem cell.

### PAGA

The trajectory inference graph was generated using PAGA, which was implemented in Scanpy (v1.6). Pruning was done by setting a minimum edge weight of 0.3. Additionally, the tree was rooted at the stem cluster.

### Histological analysis

Human proximal colonic tissue was fixed in 4% (v/v) neutral-buffered paraformaldehyde, embedded in paraffin, and cut into 10 μm sections. Immunofluorescent staining of LYZ was performed to visualize Paneth cells. Briefly, sections were incubated with primary antibody (rabbit anti-LYZ, 1:1000 dilution in PBS with 1% (w/v) BSA) (Invitrogen, Carlsbad, CA, cat. PA5-16668) overnight at 4° C followed by goat anti-rabbit Alexa fluor 594 secondary antibody (1:1000 in PBS with 1% (w/v) BSA) (Invitrogen, Carlsbad, CA, cat. A1102) incubation for 1 hr at room temperature. Subsequent section incubation with DAPI (1:1000 in PBS) (Invitrogen, Carlsbad, CA, cat. D1306) for 30 min. at room temperature was used to visualize nuclei. Images were captured using a BX53 Olympus scope (Olympus, Center Valley, PA). LYZ+ cells were enumerated in longitudinally well-oriented colonic crypts.

## Acknowledgments

This work was funded in part through Helmsley Charitable Trust (SHARE Project 2), NIDDK P01DK094779, NIDDK 1R01DK104828-01A1, NIDDK P30-DK034987, NIH T32 Translational Medicine Training Grant (T32-GM122741), and Research Fellow Award from Crohn’s and Colitis Foundation. The UNC Translational Pathology Laboratory is supported, in part, by grants from the National Cancer Institute (3P30CA016086).

## Figure Legends

**Supplemental Figure 1:**
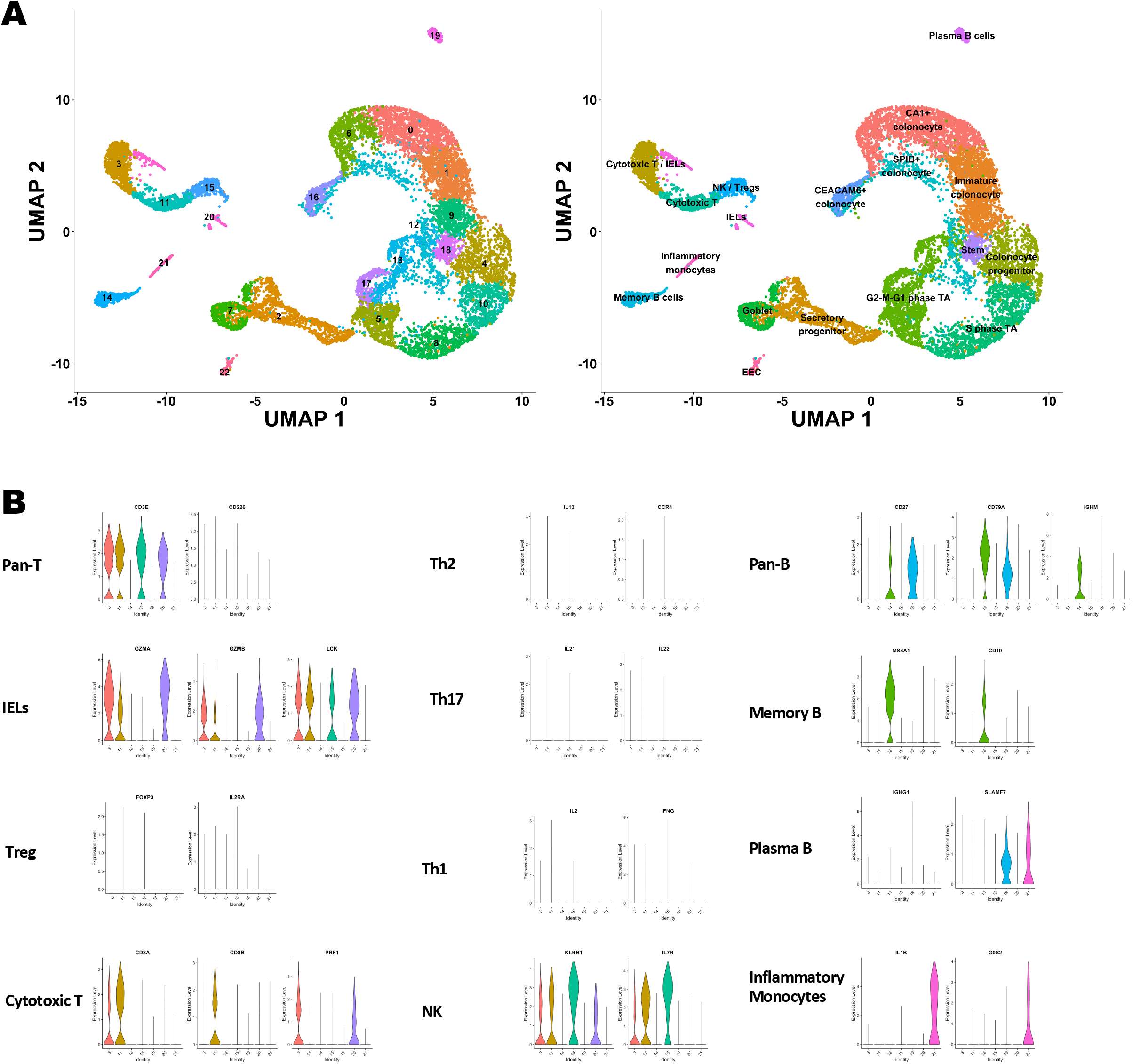
Clustering and identification of contaminating immune cells. (A) UMAP of initial clusters (left) and clusters following cell type annotation using known cell type markers and highly enriched genes (right). (B) Violin plots showing normalized expression of known markers of different immune cell types on the y-axis and the immune clusters on the x-axis. UMAP = Uniform Manifold Approximation and Projection.

**Supplemental Figure 2:**
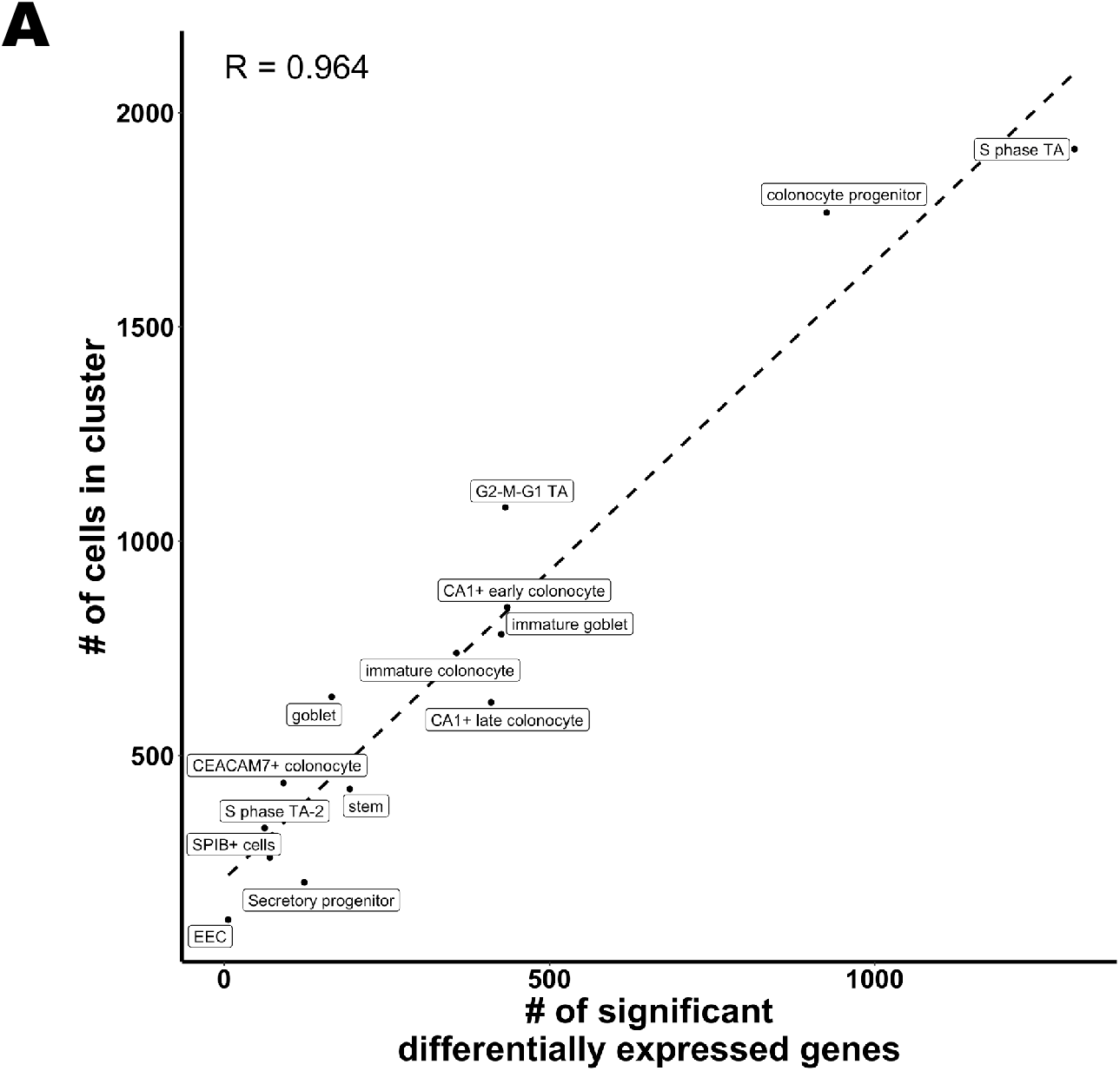
Correlation between cluster size and identification of significant differentially expressed genes. (A) The number of significant differentially expressed genes (x-axis) increases with number of cells in cluster (y-axis). Dashed line is the linear model with the corresponding R value shown in upper quadrant of plot.

**Supplemental Figure 3:**
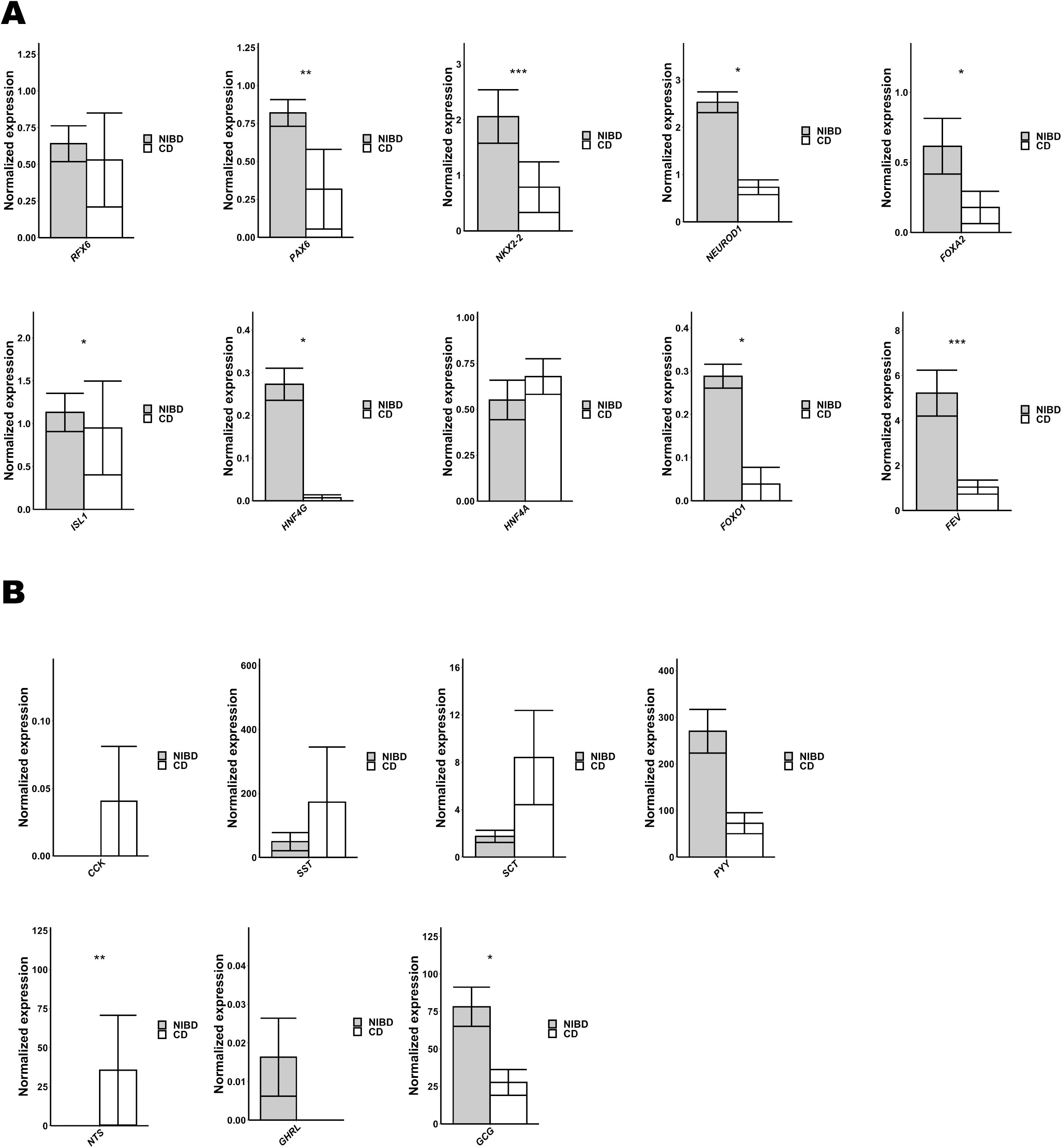
CD disrupts EEC homeostatsis. (A) Mean normalized expression across 10 transcription factors involved in EEC maturation in NIBD and CD samples. (B) Mean normalized expression across 7 EEC hormones in NIBD and CD samples. P-values were calculated using the Wilcoxon Rank Sum test (p < 0.05 = *, p < 0.01 = **, p < 0.001 = ***). NIBD = Non-inflammatory bowel disease, CD = Crohn’s disease, EEC = Enteroendocrine cells.

**Supplemental Figure 4:**
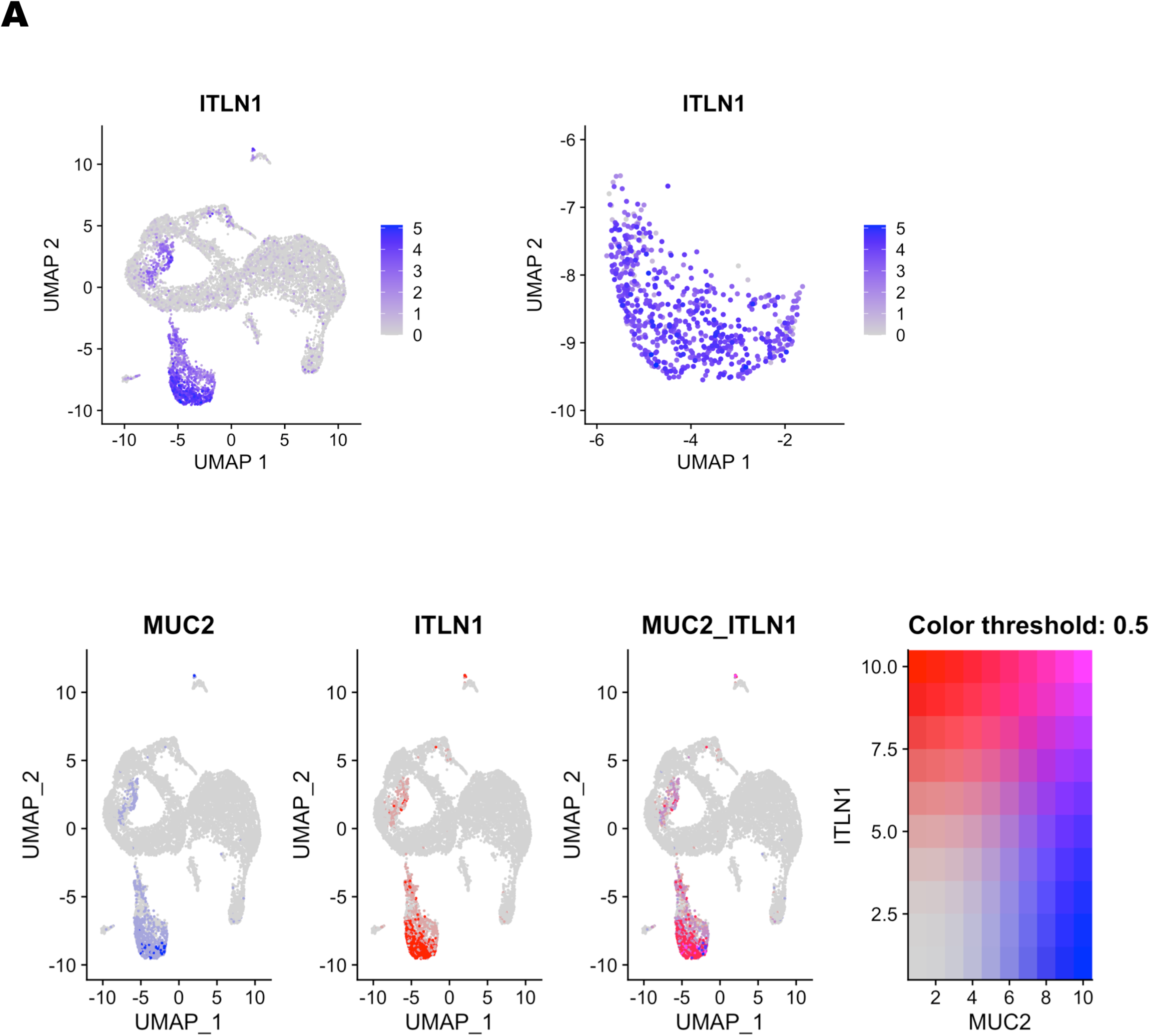
Expression of ITLN1 in mature goblet cells. (A) UMAP of all epithelial clusters (top left) or the mature goblet cluster (top right) overlain with *ITLN1* expression. UMAP of all epithelial clusters overlain with both *MUC2* (blue) and *ITLN1* (red) expression (bottom).

**Supplemental Table 1:**
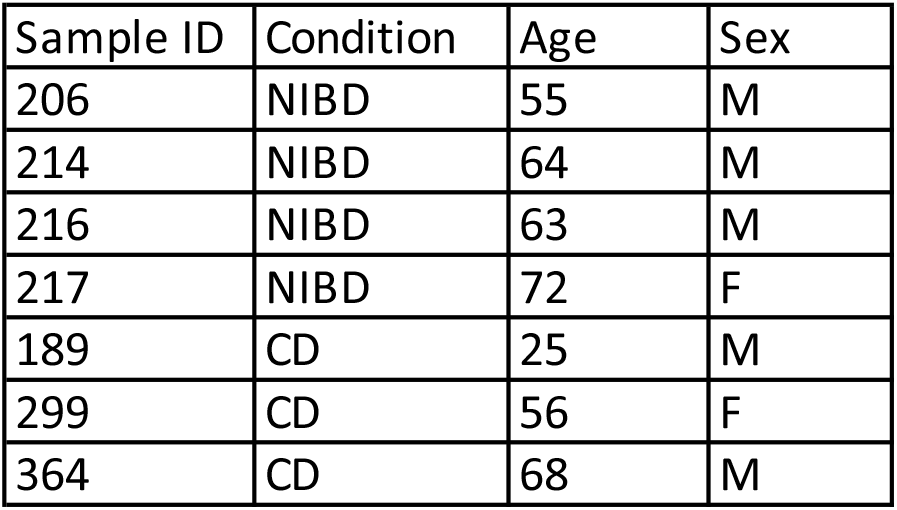
Information on the seven individuals from which the single cell tissue was biopsied.

| Sample ID | Condition | Age | Sex |
| --- | --- | --- | --- |
| 206 | NIBD | 55 | M |
| 214 | NIBD | 64 | M |
| 216 | NIBD | 63 | M |
| 217 | NIBD | 72 | F |
| 189 | CD | 25 | M |
| 299 | CD | 56 | F |
| 364 | CD | 68 | M |

**Supplemental Table 2:**
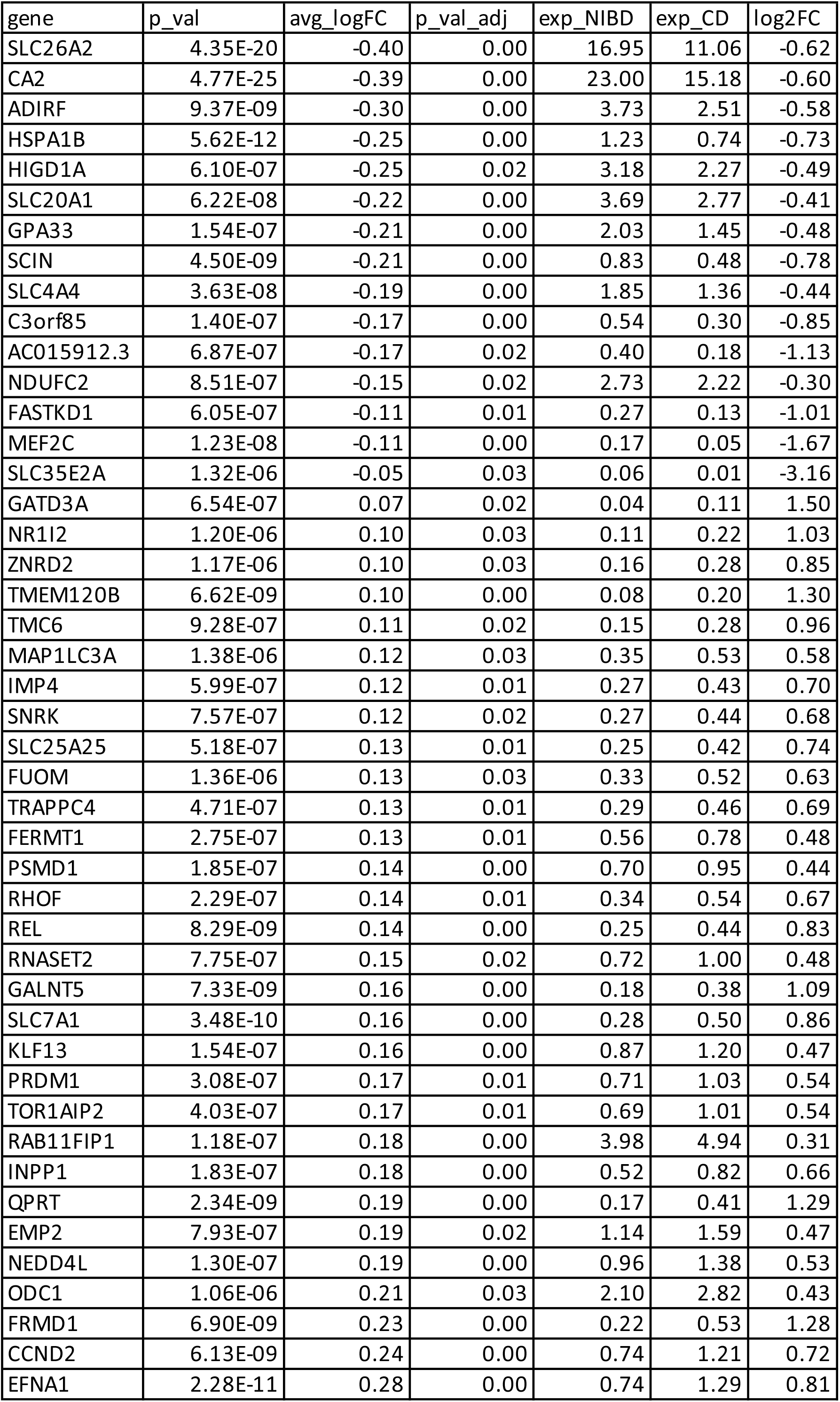
Significant (p-adjusted < 0.05) differentially expressed genes in CD compared to NIBD that are specific to *CA1*+ late colonocytes.

**Supplemental Table 3:**
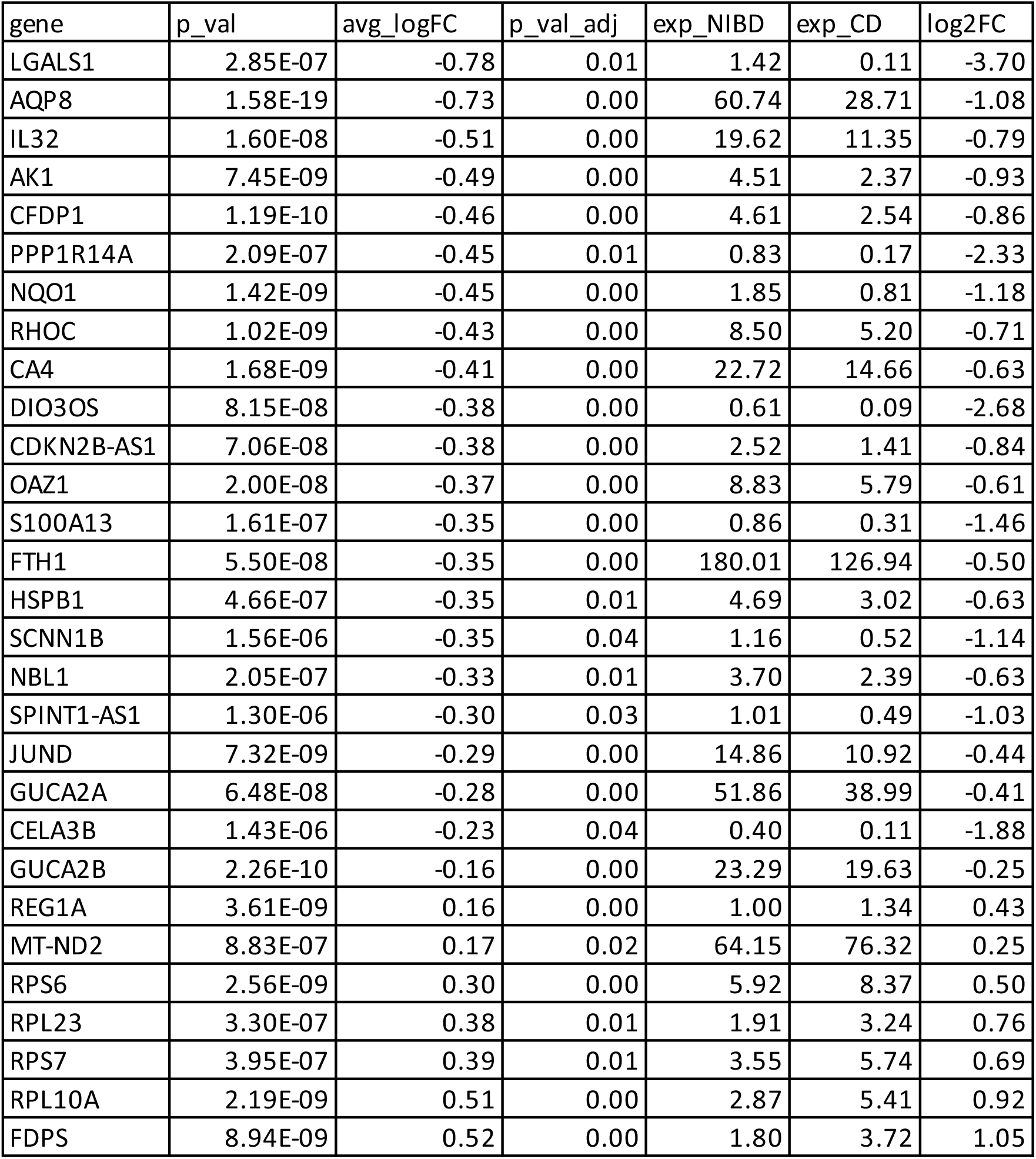
Significant (p-adjusted < 0.05) differentially expressed genes in CD compared to NIBD that are specific to *CEACAM7*+ colonocytes.

**Supplemental Table 4:** Average UMI count, detected genes, and percent of mitochondrial genes per cell across the seven samples used.

| Sample ID | Condition | Avg. UMI per cell | Avg. genes per cell | Avg. % of mitochondrial reads per cell |
| --- | --- | --- | --- | --- |
| 206 | NIBD | 21670.1 | 4153.1 | 20.1 |
| 214 | NIBD | 21562 | 4216.2 | 20.1 |
| 216 | NIBD | 23015.5 | 4187.4 | 20.8 |
| 217 | NIBD | 21538.3 | 4104.8 | 18.9 |
| 189 | CD | 19781 | 3895.8 | 20.2 |
| 299 | CD | 17240.9 | 3592.3 | 21.2 |
| 364 | CD | 20114 | 3991.2 | 18.7 |

